# Lymphatic dysfunction in lupus contributes to cutaneous photosensitivity and lymph node B cell responses

**DOI:** 10.1101/2022.06.13.495930

**Authors:** Mir J. Howlader, William G. Ambler, Madhavi Latha S. Chalasani, Aahna Rathod, Ethan S. Seltzer, Ji Hyun Sim, Jinyeon Shin, Noa Schwartz, William D Shipman, Dragos C. Dasoveanu, Camila B. Carballo, Ecem Sevim, Salma Siddique, Yurii Chinenov, Scott A. Rodeo, Doruk Erkan, Raghu P. Kataru, Babak Mehrara, Theresa T. Lu

## Abstract

Patients with systemic lupus erythematosus (SLE) are photosensitive, developing skin inflammation with even ambient ultraviolet radiation (UVR), and this cutaneous photosensitivity can be associated with UVR-induced flares of systemic disease, which can involve increased autoantibodies and further end organ injury. Mechanistic insight into the link between the skin responses and autoimmunity is limited. Signals from skin are transmitted directly to the immune system via lymphatic vessels, and here we show evidence for potentiation of UVR-induced lymphatic flow dysfunction in SLE patients and murine models. Improving lymphatic flow by manual lymphatic drainage (MLD) or with a transgenic model with increased lymphatic vessels reduces both cutaneous inflammation and lymph node B and T cell responses, and long term MLD reduces splenomegaly and titers of a number of autoantibodies. Mechanistically, improved flow restrains B cell responsesS in part by stimulating a lymph node fibroblastic reticular cell-monocyte axis. Our results point to lymphatic modulation of lymph node stromal function as a link between photosensitive skin responses and autoimmunity and as a therapeutic target in lupus, provide insight into mechanisms by which the skin state regulates draining lymph node function, and suggest the possibility of MLD as an accessible and cost-effective adjunct to add to ongoing medical therapies for lupus and related diseases.

## Introduction

Photosensitivity, a cutaneous sensitivity to ultraviolet radiation (UVR), affects the majority of systemic lupus erythematosus (SLE) patients, but, in addition to inflammatory skin lesions, UVR exposure can trigger systemic disease flares in both patients and murine models that can include increased autoantibody levels and further end organ injury (1–5). Currently, medications used for photosensitive skin responses include topical steroids, topical calcineurin inhibitors, and antimalarials such as hydroxychloroquine. More importantly, lupus patients are advised to reduce UVR exposure by avoiding the sun, wearing protective clothing, and wearing sunscreen to prevent photosensitive skin responses and their sequelae (6–8). Photosensitive skin inflammation, the accompanying risk of systemic disease flares, and the lifestyle modifications needed to prevent these all contribute to disrupting patients’ quality of life (9–11). Advances have begun to delineate the mechanisms that contribute to photosensitivity (12–14). However, the link between UVR-induced skin inflammation and increased autoantibody titers in these photosensitive patients remains poorly understood.

Skin communicates with lymphoid tissues via lymphatic vessels which transport cells and interstitial fluid to skin-draining lymph nodes where immune responses occur and can be regulated. This lymphatic transport serves both to clear fluid and inflammation from the skin and to deliver antigens, antigen-presenting cells, and mediators that impact lymph node function (15, 16). Reduced lymphatic flow results in exacerbated skin inflammation and, over time, can lead to the development of autoantibodies (17, 18). This latter scenario is not well understood mechanistically but the combination of skin inflammation and autoimmunity is reminiscent of SLE, and raises the possibility of lymphatic dysfunction in SLE, which has been described anecdotally (19–22).

Lymphatic flow brings both dendritic cells and lymph fluid to the subcapsular sinus of the draining lymph node, at which point they take divergent paths. Dendritic cells migrating from the skin actively leave the sinus to enter the nodal parenchyma where T and B cells are located (23). Lymph fluid containing soluble molecules, on the other hand, flows into the conduit system (24). The conduits consist of a central collagen core ensheathed by fibroblastic reticular cells (FRCs), and the potential space between the collagen core and the FRCs allows for flow of lymph fluid throughout the lymph node until the lymph leaves the node mainly via efferent lymphatic flow. Comprising the wall of the conduit system, then, FRCs are among the major first sensors of signals flowing from the skin. FRCs, in turn, play critical roles regulating T and B cell responses (25–27), and we recently showed that immunization of healthy mice upregulated FRC-derived CCL2 which promoted CCR2+ Ly6C+ monocyte expression of reactive oxygen species (ROS) to limit plasmablast survival (28). Lymphatic input from skin to lymph nodes that alters FRC phenotype, then, could potentially impact lymph node B cell responses.

In this study, we examined for lymphatic dysfunction in both human SLE and murine SLE models and sought to understand the consequences on both skin and lymph node function. We show evidence for potentiation of UVR-induced flow dysfunction in human SLE and in multiple photosensitive SLE models. Improving lymphatic flow by manual lymphatic drainage (MLD) or with a transgenic model with increased lymphatic flow reduced both UVR-induced skin inflammation, draining lymph node B and T cell responses, and MLD over a prolonged period of time reduced splenomegaly and titers of a number of autoantibodies. We further show that improving lymphatic flow upregulates FRC CCL2, and that depleting monocytes limits the flow-induced reduction in plasmablasts. These results suggest that SLE skin is primed for UVR-induced lymphatic flow dysfunction, and the dysfunction contributes to both cutaneous photosensitive responses and, by modulating a FRC-monocyte axis, lymph node B cell activity in disease. This scenario suggests that improving lymphatic flow and its consequences on the lymph node stromal microenvironment may be therapeutically useful in SLE.

## Results

### Evidence of dysfunctional dermal lymph flow in SLE patients and mouse models

We assessed for evidence of lymphatic flow alterations in the skin of SLE patients and SLE mouse models. In human skin, greater lumenal area of lymphatic vessels can be reflective of distention from reduced lymphatic flow (29–31). We examined biopsies that we had previously obtained from the sun-exposed forearm skin of patients with SLE and with persistently positive antiphospholipid antibodies (APL), a condition that can overlap with SLE. These biopsies were taken from sites of livedo reticularis, a lacy pattern of prominent veins on otherwise normal-appearing skin that is not considered to be related to photosensitivity and that affects both SLE and APL patients (32). We compared the samples from SLE patients (some of whom also had APL) to that of APL patients without SLE. SLE skin showed dilated lymphatic vessels when compared to APL-only skin **(Figure 1A-B**). There was no change in the density of lymphatic vessels **(Figure 1C).** While these results are in the context of livedo skin, the differences between SLE and non-SLE patients suggested that there is dysfunctional lymphatic flow in human SLE skin that is exposed to UVR.

**Figure 1.**
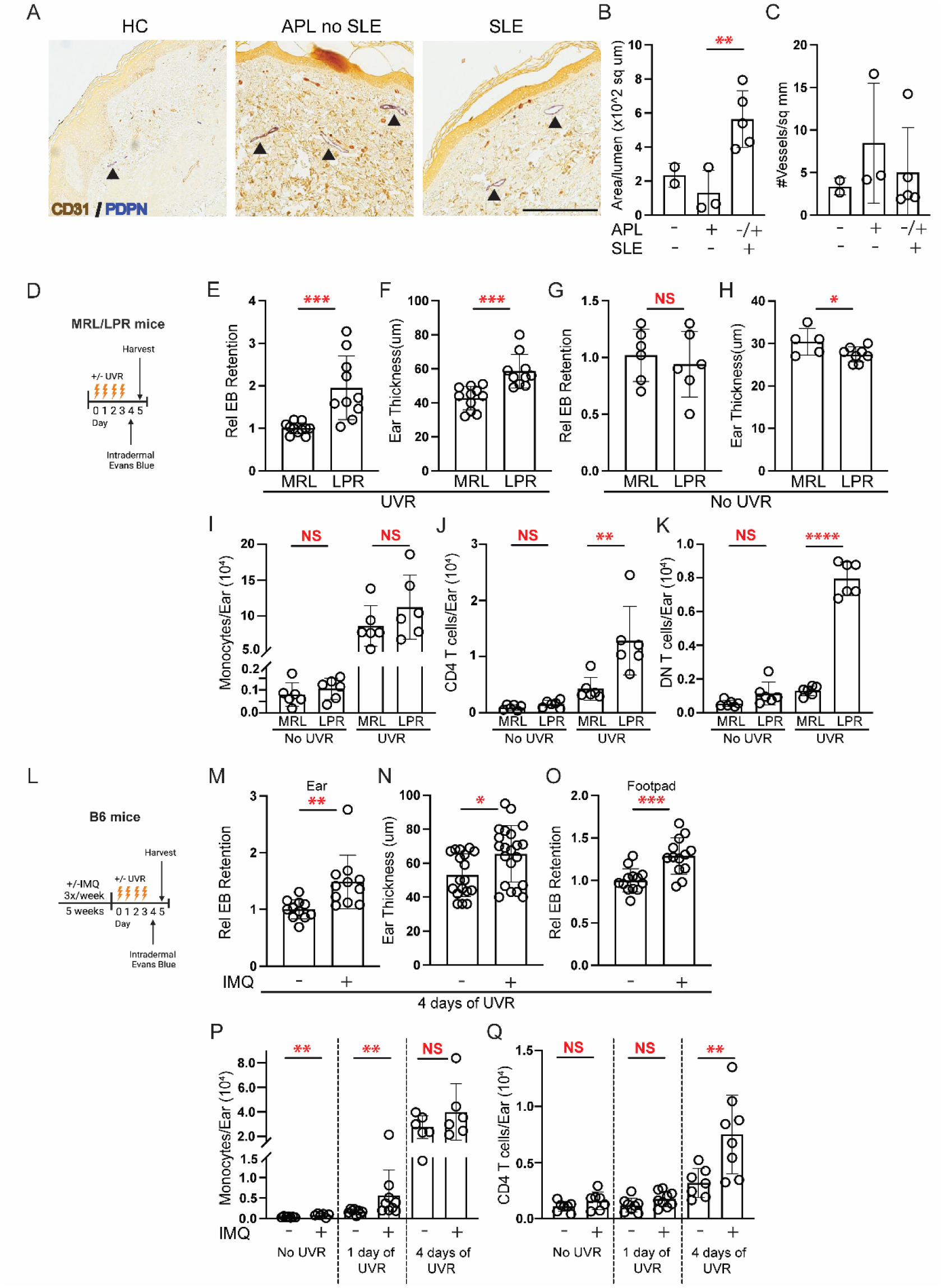
Patients with SLE and murine SLE models show evidence of cutaneous lymphatic dysfunction. (A-C) Punch biopsies of forearm skin from healthy control (HC), positive antiphospholipid antibodies (APL) without SLE, and SLE subjects with or without APL were stained for CD31+PDPN+ lymphatic vessels. (A) Representative photomicrographs. Arrowheads point to lymphatic vessels. (B) Lumenal area per vessel. Each symbol is an individual subject. (C) Number of lymphatic vessels per tissue area. (D-H) LPR mice and MRL controls were treated with ultraviolet radiation (UVR) for 4 consecutive days, injected in ear pinna with Evans blue dye (EB) 1 day after final UVR dose, and ear harvested to assess EB content 1 day later, as in (D). (E) EB retention and (F) ear swelling was quantified after UVR or (G,H) with no UVR. (I-K) Ears were examined by flow cytometry at 24 hours after final dose of UVR. (I) Monocyte, (J) TCRαβ CD4+ T cell, and (K) TCRab+ CD3+ CD4-CD8-“double negative” (DN) T cell numbers. (L-O) B6 mice received IMQ on the right ear and exposed to UVR, and EB retention after intradermal injection of left ear or footpad was compared to responses in vehicle-treated control mice, as in (L). (M) EB retention in ear. (N) Ear swelling. (O) EB retention in footpad. (P-Q) Left ear was collected 24 hours after 1 or 4 doses of UVR. (P) Monocyte and (Q) TCRαβ CD4 T cell numbers. Each symbol represents one mouse; n = 2 to 21 per condition; data are from 2 (I, J, K), 3 (M, O), 4 (G, H), 6 (E, F), 10 (N), and 11 (P, Q) independent experiments. Normality was assessed using the Shapiro-Wilk test. If normal, unpaired t test was used. If data were not normal, Mann-Whitney U test was used. ***P<0.001, **P<0.01, *P<0.05, NS=not significant. Error bars represent SD.

We examined lymphatic flow in lupus model mice by assaying for retention in the skin of intradermally injected Evans blue dye (33) after UVR treatment. We focused on ear skin as it has little fur, allowing the skin to be directly exposed to UVR. We used two photosensitive lupus models, the MRL-*Fas*^lpr^ (LPR) spontaneous lupus model (34–36) and the imiquimod (IMQ) lupus model that is induced by chronic epicutaneous IMQ application (37). The LPR mice and their MLR-MpJ (MRL) controls were treated with 1000J UVB/m2/day of UVR for 4 days prior to Evans blue dye injection in the ear skin (**Figure 1D**). LPR mice showed increased Evans blue dye retention in the ear when compared to MRL mice **(Figure 1E)**, suggesting reduced flow out of the ear skin. As expected for the photosensitive LPR mice, there was greater UVR-induced skin swelling in LPR mice when compared to control MRL mice **(Figure 1F).** The greater Evans blue dye retention in LPR mice was specific to UVR exposure, as LPR mice showed no increases in Evans blue dye retention or ear thickness at baseline without UVR (**Figure 1G-H)**.

The UVR-induced lymphatic alterations and ear swelling were accompanied by the accumulation of inflammatory cells. We had previously shown that inflammatory Ly6C^hi^ monocytes accumulated after one day of UVR in LPR mice (35). After 4 days of UVR, monocytes numbers accumulated in greater numbers in both MRL and LPR mice treated with UVR at comparable levels **(Figure 1I)**. However, TCRαβ+ CD4 T cells and TCRαβ+CD3+CD4-CD8-“double negative” (DN) T cells that are characteristic of LPR mice (38) both accumulated in greater numbers in LPR mice compared to MRL mice at this time point **(Figure 1J-K)**. Neutrophil, CD8 T cells, and TCRγδ T cells showed no changes between LPR and MRL mice **(Supplemental Figure 1A-C)**. Together with the Evans blue dye experiments, these results indicated that lymphatic flow dysfunction accompanies the increased inflammation in the skin (as indicated by the greater edema and T cell infiltrate) in UVR-treated LPR mice. These results suggested the possibility that lymphatic flow dysfunction, by failing to remove fluid and inflammatory mediators from the skin, was contributing to the increased skin inflammation of the lupus model mice.

We examined the IMQ model (37), inducing this model in B6 mice by applying IMQ, a TLR7 agonist, to the skin for 4-5 weeks, yielding B6-IMQ mice **(Figure 1L)**. IMQ was applied on the right ear only, and the skin on the rest of the body, including the left ear, was considered “non-lesional” skin and reflective of systemic disease. We have shown previously that the “non-lesional” left ear in IMQ mice is similar to non-lesional skin in human lupus in expressing an interferon signature whereas the right ear has a less robust interferon signature (39). Additionally, the right ear, even without UVR exposure, showed upregulation of apoptotic pathways (**Supplemental Figure 2A**), suggesting that there was tissue damage from repeated local treatment of IMQ. To assess effects of UVR exposure without confounding results from direct IMQ treatment, we focused on the left, non-lesional ear. The left ear showed increased Evans blue dye retention with increased skin swelling upon UVR exposure when compared to vehicle-painted controls **(Figure 1M-N)**. Similarly, the footpad of B6-IMQ mice showed increased Evans blue dye retention when compared to B6 mice (**Figure 1O**), suggesting that the dermal lymphatic dysfunction in B6-IMQ mice affected non-lesional skin throughout the body and that the left ear results were not reflective of direct exposure to IMQ that had transferred from the right ear. The right ear also showed increased Evans blue dye retention and increased ear thickness (**Supplemental Figure 2B-C**).

Characterization of the inflammatory infiltrate in B6-IMQ mice showed a higher baseline monocyte number in B6-IMQ compared to B6 mice, and UVR exposure for 1 day showed greater monocyte accumulation in B6-IMQ mice (**Figure 1P**). Monocyte accumulation continued to increase with additional days of UVR exposure, and were equally high in B6-IMQ and B6 ears after 4 days of UVR (**Figure 1P**). In contrast, CD4 T cell number accumulation was limited until after 4 days of UVR, when B6-IMQ showed higher numbers of CD4 T cells than B6 mice (**Figure 1Q**). Neutrophils, TCRαβ CD8 T cells, and TCRγδ T cells showed no changes between IMQ and control mice after 4 days of UVR exposure (**Supplemental Figure 1D-F**). Similar to LPR mice, then, B6-IMQ mice showed that UVR-induced skin inflammation after 4 days of treatment included lymphatic dysfunction along with tissue swelling and T cell accumulation. Together, our findings in two distinct SLE models suggested that photosensitive skin is characterized in part by UVR-induced lymphatic dysfunction, supporting the evidence of lymphatic dysfunction in sun-exposed skin in human SLE.

### Improving lymphatic flow reduces photosensitivity of skin in SLE models

We asked about the contribution of reduced lymphatic flow to cutaneous photosensitive responses in the SLE models by using two different approaches to improve lymphatic flow. One approach was through manual lymphatic drainage (MLD), a technique used by physical therapists to reduce swelling in patients with acquired or congenital lymphedema (40, 41). We administered MLD targeting the left ear once a day during the course of UVR exposure **(Figure 2A-B).** This treatment decreased Evans blue retention in LPR mice relative to handling controls **(Figure 2C and Supplemental Figure 3A)**, suggesting that MLD was successful in improving lymphatic flow. MLD also ameliorated UVR-induced skin inflammation, with reductions in ear swelling **(Figure 2D-E),** CD4 T cell numbers and interferon gamma (IFNg) expression **(Figure 2F-H)**, and DN T cell numbers and IFNg expression **(Figure 2I-J)**. The proportion of regulatory T cells (Treg) and T helper 17 cells (Th17) within the CD4 T cell population, the proportion of IL-17-expressing DN cells, and the number of CD8 T cells did not change with MLD (**Supplemental Figure 3B-E**). Monocyte and neutrophil numbers also remained unchanged (**Supplemental Figure 3F-G).** These results suggested that improving lymphatic flow with MLD was able to both reduce ear swelling and T cell accumulation and activity in UVR-treated LPR mice.

**Figure 2.**
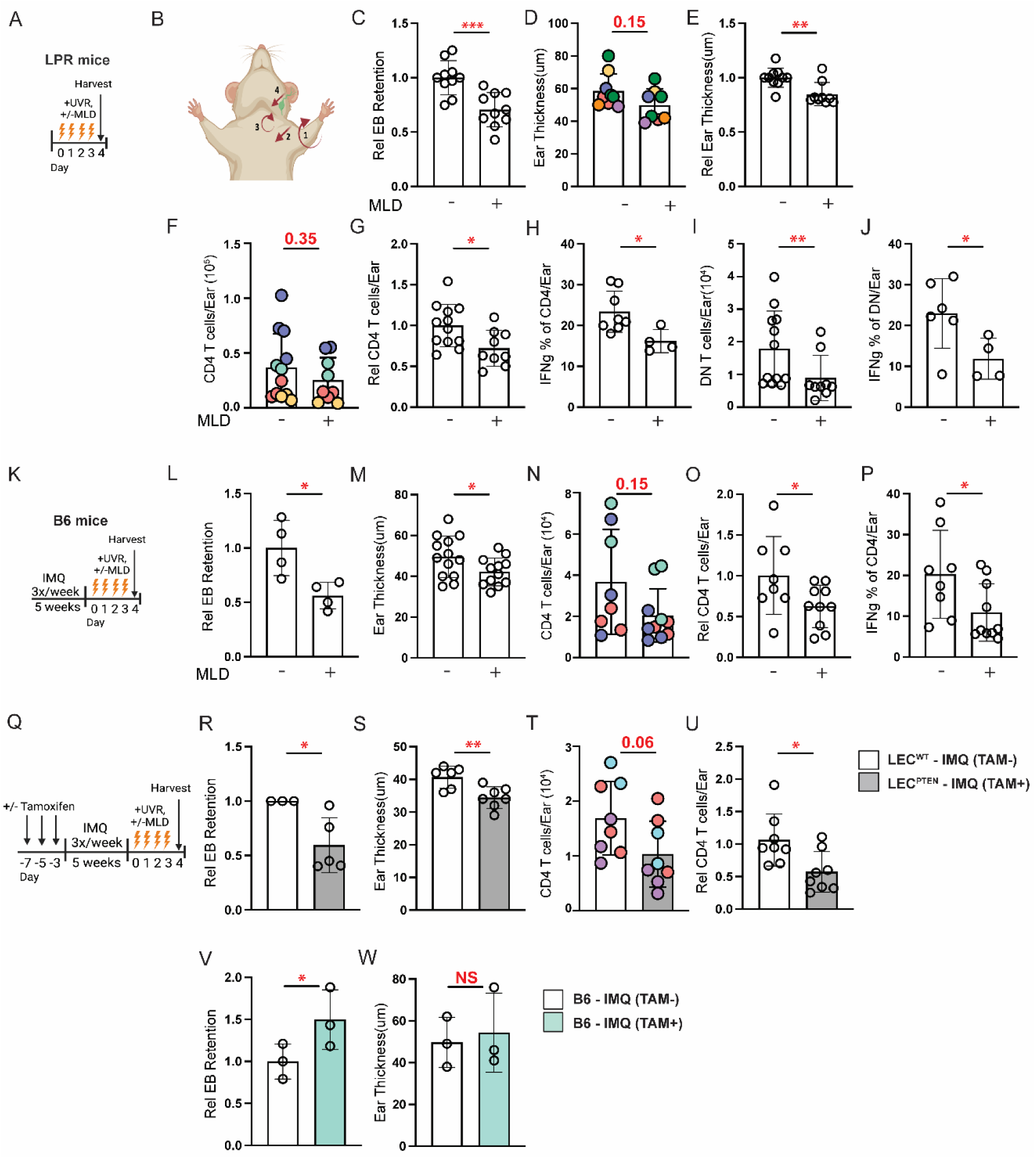
Improving lymphatic flow reduces cutaneous photosensitive responses in SLE models. (A-J) LPR mice and (K-P) B6-IMQ mice were treated with UVR and manual lymphatic drainage (MLD) targeting the left ear or were control handled. Left ear was then examined. (A, K) Experimental design. (B) Illustration of MLD technique. Please see Methods for details. (C, L) EB retention and (D, E, M) ear thickness, absolute (D,M) or normalized (E). (F, N) TCRαβ CD4 T cell numbers, (G,O) normalized to controls. (H, P) Percentage of CD4 T cells that express IFNg. (I) DN T cell numbers and (J) percentage that express IFNg. (Q-U) Flt4Cre^ERT2^ PTEN^fl/fl^ mice treated with tamoxifen (LEC^PTEN^) or without (LEC^WT^) were treated with IMQ on right ear and UVR before left ear skin assessment, as in (Q). (R) EB retention, (S) skin thickness, (T) TCRαβ CD4 T cell numbers, and (U) IFNg+ percentage. (V, W) Non-transgenic B6 mice were treated as described in (Q). (V) EB retention and (W) ear thickness. Each symbol represents one mouse; n= 3 to 13 per condition; data are from 2 (H, J, L, V, W), 3 (N, O, P, R, T, U), 4 (F, G, I, S), 6 (C), 7 (D, E), and 9 (M) independent experiments. Normality was assessed using the Shapiro-Wilk test. If normal, unpaired t test was used. If data were not normal, Mann-Whitney U test was used. ***P<0.001, **P<0.01, *P<0.05, NS=not significant. Error bars represent SD.

MLD had similar effects in B6-IMQ mice **(Figure 2K-P)**. Evans blue dye retention was reduced **(Figure 2L)**, suggesting improved lymphatic flow, and this was accompanied by reduced ear swelling **(Figure 2M)**. Both total CD4 T cell numbers and IFNg-expressing Th1 proportion were also decreased with MLD **(Figure 2N-P)**. Similar to LPR mice, MLD had no effect on Treg, Th17, CD8 T cell, monocyte, or neutrophil accumulation **(Supplemental Figure 3H-L)**. Our results together suggested that improving lymphatic flow with MLD ameliorated UVR-induced cutaneous inflammation in both the LPR and IMQ SLE models.

Our second approach to improve lymph flow was by using transgenic Flt4Cre^ERT2^PTEN^fl/fl^ mice (42). When treated with tamoxifen, these mice have lymphatic endothelial cell (LEC)-specific deletion of PTEN, an antagonist of critical VEGFC/VEGRF3 signaling. This results in expansion of functional lymphatic vessels, improved lymphatic flow from skin, and reduced UVR-induced skin inflammation in healthy (ie non-lupus) mice (42). We confirmed that tamoxifen treatment of Flt4Cre^ERT2^PTEN^fl/fl^ mice specifically deleted PTEN from LECs (**Supplemental Figure 4A**). We designated the tamoxifen-treated mice as “LEC^PTEN^ mice” and non-tamoxifen-treated mice as “LEC^WT^ mice”, and induced the IMQ model in them to generate LEC^PTEN^-IMQ and LEC^WT^-IMQ mice (**Figure 2Q**). Reduced Evans blue dye retention in the left ear of LEC^PTEN^-IMQ mice compared to LEC^WT^-IMQ controls confirmed improved lymphatic drainage (**Figure 2R**), and this was associated with reduced UVR-induced ear swelling **(Figure 2S),** suggesting that improving lymphatic flow reduced the ear swelling. Similar to MLD, the genetic approach to improving lymphatic flow reduced CD4 T cell numbers in LEC^PTEN^-IMQ mice compared to LEC^WT^-IMQ mice (**Figure 2T-U**). Notably, tamoxifen treatment of non-transgenic B6 mice did not reduce Evans blue dye retention or ear swelling **(Figure 2V-W),** suggesting that effects in LEC^PTEN^-IMQ mice were attributable to PTEN deletion rather than to tamoxifen treatment. Treg, CD8 T cell, monocyte, and neutrophil numbers remained unchanged (**Supplemental Figure 3M-P**). The right ear of LEC^PTEN^-IMQ mice also showed reduced Evans blue dye retention and ear swelling when compared to LEC^WT^-IMQ mice and no changes in monocyte and neutrophil numbers (**Supplemental Figure 4B-E**). As with the left ear, B6-IMQ mice showed no reduction in Evans blue dye retention or ear thickness with tamoxifen (**Supplemental Figure 4F-G**). Consistent with prior reports that improved lymphatic drainage can reduce cutaneous inflammation (33, 42, 43), our results from using both an acute physical approach in two lupus models and a long-term genetic approach showed that improving lymphatic flow from skin reduces cutaneous UVR-induced inflammation in SLE model mice, suggesting that the lymphatic dysfunction contributes to photosensitive skin responses in SLE.

### Improving lymphatic flow reduces draining lymph node B cell responses in SLE models

We asked whether lymphatic flow alterations in SLE models contributed to modulating immune activity in downstream lymph nodes. Improving lymphatic flow with MLD over 4 days in LPR mice during UVR exposure did not affect overall lymph node cellularity or overall B cell numbers in draining auricular nodes (**Figure 3A-B**), but did reduce germinal center B cell and plasmablast numbers (**Figure 3C-D**). CD4 T cell numbers were also reduced (**Figure 3E-F**), as was the frequency of TFH cells (**Supplemental Figure 5A**), which could have contributed to reduced germinal center B cell responses. Th1 (**Figure 3G**), Tregs, and Th17 frequencies were unchanged (**Supplemental Figure 5B-C**). CD8 (**Figure 3H**), and DN T cell (**Figure 3I**) numbers were unchanged, as was the expression of IL-17 by DN cells (**Supplemental Figure 5D**). These data suggested that improving lymphatic flow with MLD in LPR mice reduces draining lymph node germinal center and plasmablast responses along with overall CD4 T cell numbers and the proportion of TFH cells.

**Figure 3.**
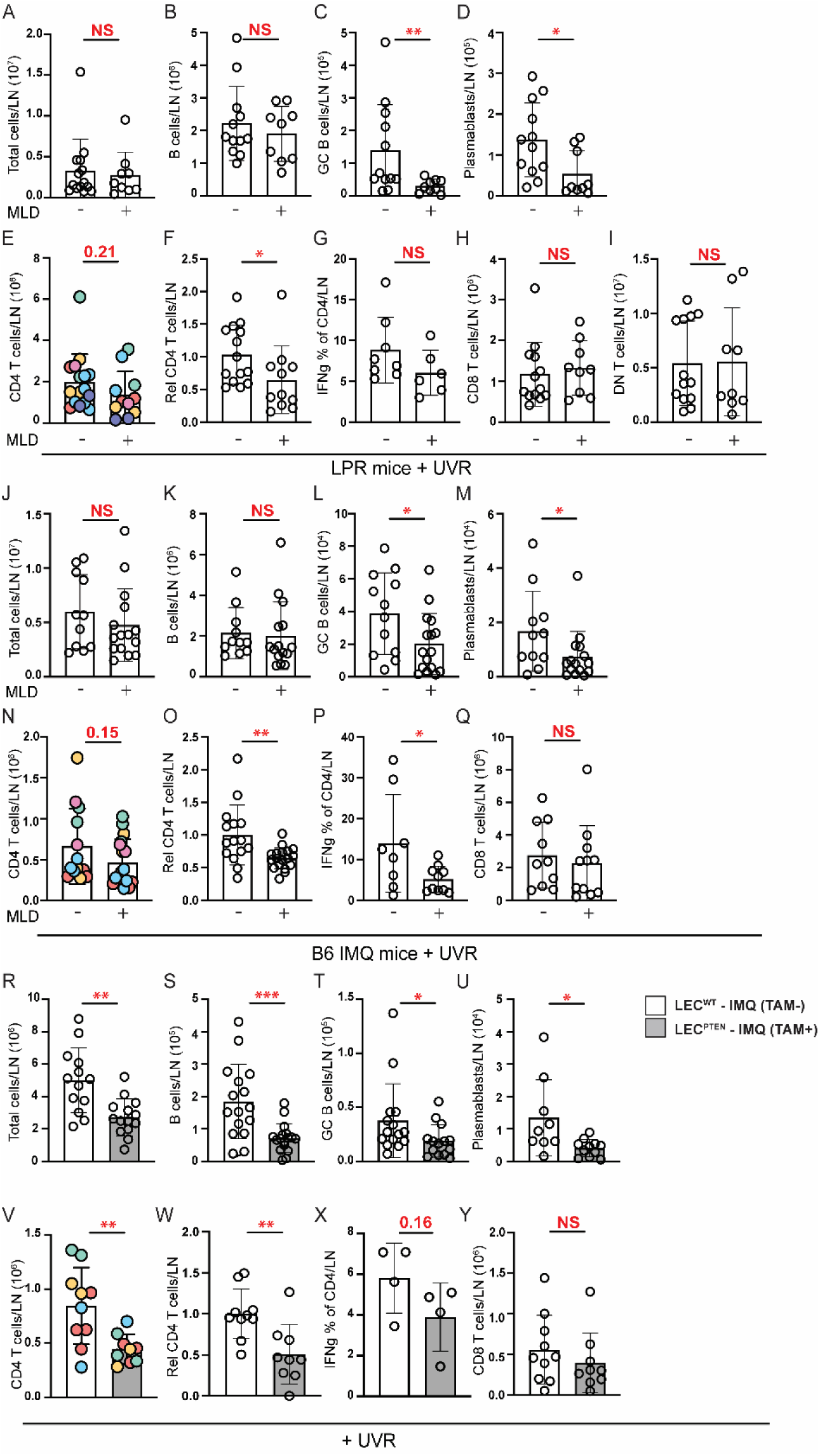
Improving lymphatic flow reduces draining lymph node B and T cell responses in SLE models. (A-Q) Left auricular lymph nodes of (A-I) LPR, (J-Q) B6-IMQ, and (R-X) LEC^PTEN^-IMQ and LEC^WT^-IMQ mice that were treated as in Figure 2A, K, and Q were examined. (A, J, R) Lymph node cellularity. (B, K, S) B cell, (C, L, T) germinal center B cell, (D, M, U) plasmablast, and (E, N, V) TCRαβ CD4 T cell numbers. (F, O, W) CD4 T cell numbers normalized to control. (G, P, X) Percentage of CD4 T cells that express IFNg. (H, Q, Y) CD8 and (I) DN T cell numbers. Each symbol represents one mouse; n= 4 to 17 per condition; data are from 2 (X), 3 (G, P, Q), 4 (A-C, H-K, M, V, W, Y), 5 (D, N, O), 6 (E, F, L, R, U), and 8 (S, T) independent experiments. Normality was assessed using the Shapiro-Wilk test. If normal, unpaired t test was used. If data were not normal, Mann-Whitney U test was used. ***P<0.001, **P<0.01, *P<0.05, NS=not significant. Error bars represent SD.

Similar to LPR mice, MLD in B6-IMQ mice showed no effects on total LN cellularity or on overall B cells (**Fig 3J-K)** but did reduce germinal center B cell, plasmablast, and CD4 T cell numbers (**Fig 3L-O)**. Note that the reduction in germinal center (2.2 fold) and plasmablast (2.4 fold) numbers reflects a partial reduction rather than normalization to a non-lupus state, as UVR-treated B6 mice have very few germinal center cells and plasmablasts (1002+/-859 and 312+/-220 per lymph node, respectively). Unlike in LPR mice, MLD in B6-IMQ mice did not reduce TFH cells **(Supplemental Figure 5E)** but did reduce the TH1 frequency **(Figure 3P).** Tregs and Th17 cell frequencies **(Supplemental Figure 5F-G)** and CD8 T cell numbers **(Figure 3Q)** were unchanged.

These data together showed that MLD in LPR and IMQ models reduced B cell responses and CD4 T cell numbers and differentiation, thus suggesting that even a short period of improved lymphatic flow from skin can reduce B and CD4 T cell responses.

In contrast to LPR and B6-IMQ mice treated with MLD, LEC^PTEN^-IMQ mice showed decreases in lymph node cellularity and overall B cells when compared to LEC^WT^-IMQ mice **(Figure 3R-S).** As with LPR and B6-IMQ mice, germinal center B cell, plasmablast, and CD4 T cell numbers were reduced (**Figure 3T-W).** Th1 **(Figure 3X)**, TFH, and Treg frequencies **(Supplemental Figure 5H-I)** and CD8 T cell numbers **(Figure 3Y)** were unchanged. These data suggest that improving lymph flow genetically and before induction of lupus disease activity reduced overall lymph node cellularity, and, similar to short term MLD, preferentially reduced B cell and CD4 responses.

### Longer duration of improved lymphatic flow reduces systemic disease activity in SLE mice

While our MLD was targeted to the left ear and draining left auricular lymph node, reduced inflammation at the site of greatest UVR exposure (left ear versus the fur-covered back, for example) and its draining lymph node could potentially have a systemic effect, as soluble mediators including antigen and inflammatory cytokines can travel from the skin to draining lymph node and then, via efferent flow from the lymph node, out to the systemic circulation to the spleen and end organs (12). Also, reduced B and T cell responses in the draining lymph node would lead to fewer cells and autoantibodies leaving the lymph node to reach the circulation and spleen and other tissues to carry out inflammatory, effector, or memory functions. Furthermore, some inflammatory cells can also leave the skin directly into the blood circulation by reverse transmigration from tissue into blood vessels, as has been shown for neutrophils in UVR-treated skin (44).

We thus asked if improving lymphatic flow in the LEC^PTEN^-IMQ mice or by local MLD can affect parameters of systemic disease activity. We first looked at the splenomegaly that characterizes both the IMQ and LPR models (37, 45). Spleens do not have afferent lymphatics (46) and so splenic changes are considered to reflect systemic changes. LEC^PTEN^-IMQ mice had reduced splenic weight compared to LEC^WT^-IMQ mice (**Figure 4A**), suggesting reduction in systemic disease activity with improved lymphatic flow. To determine if MLD could also affect systemic disease activity, we treated LPR mice with MLD for 4-5 weeks, a duration that would allow indirect splenic changes to occur and for turnover of pre-MLD autoantibodies (47). MLD-treated LPR mice showed reduced splenic weight compared to mice that did not receive MLD, suggesting that longer term MLD can reduce systemic disease (**Figure 4B**). We further measured levels of serum autoantibodies after 4-5 weeks of UVR with and without MLD. While high titer anti-DNA and ribonucleoprotein (RNP) antibodies remained unchanged, antibodies against complement pathway components (C5, C9, Factor B, Factor I, Factor P), histone 2B, and beta2 glycoprotein were reduced (**Fig 4C,D**). Interestingly, LPR mice also expressed autoantibodies found in dermatomyositis patients (TIF1gamma, Jo1, PL-7, SAE1/SAE2, MDA5, NXP2, mi-2, PM/Scl100) who also demonstrate photosensitivity (48), and some of these (Jo-1,NXP2, PM/Scl100) were also reduced with MLD (**Fig 4C,D**). The reduced splenomegaly and autoantibody levels suggest that longer term MLD can reduce systemic disease activity.

**Figure 4.**
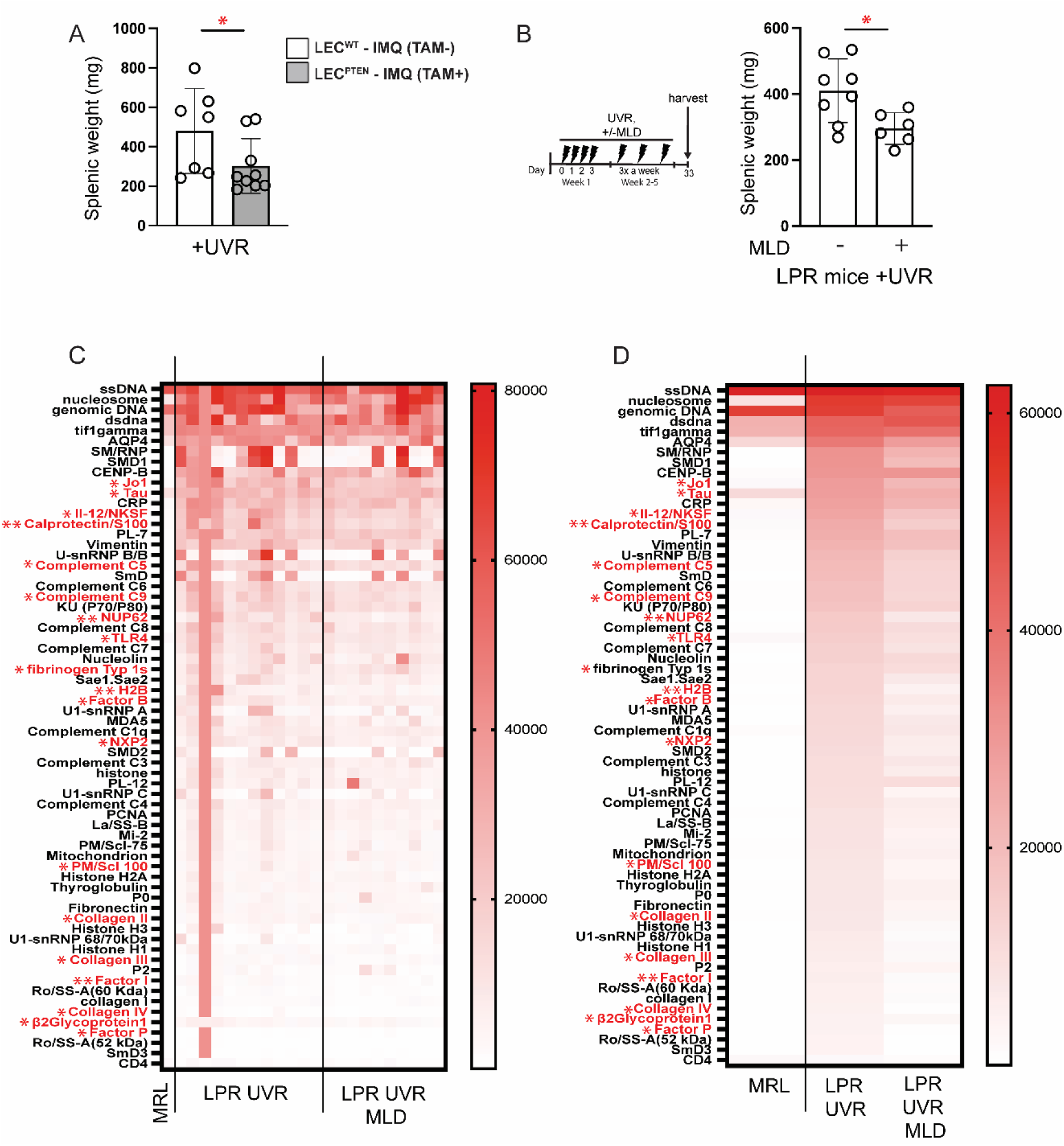
Improving lymphatic flow long term reduces systemic disease activity. Splenic weight of LEC^PTEN^-IMQ and LEC^WT^-IMQ mice treated with UVR for 4 days. (A) Splenic weight of LPR mice treated with UVR and MLD concurrently or control handled for 5 weeks. (C,D) Heatmap of normalized signal intensity (NSI) from autoantigen microarray panel for IgG of MRL mice, LPR mice treated with UVR, and LPR mice treated with both UVR and MLD for 4-5 weeks. (C) Each column represents one mouse and autoantibodies with significant differences between LPR UV and LPR UV+MLD are labelled in red. (D) Each column represents average NSI of all mice serum samples. Each symbol represents one mouse; n= 1 to 12 per condition; data are from 1 (B), 2 (C, D), and 6 (A) independent experiments. Normality was assessed using the Shapiro-Wilk test. If normal, unpaired t test was used. If data were not normal, Mann-Whitney U test was used. ***P<0.001, **P<0.01, *P<0.05, NS=not significant. Error bars represent SD.

### Improving lymphatic flow increases lymph node FRC CCL2, monocyte ROS generation, and restrains plasmablast responses in a monocyte-dependent manner

To understand how improving lymphatic flow limits lymph node B cell responses, we considered that FRCs are among the initial sensors of lymph fluid within lymph nodes and asked if improving lymphatic flow could drive an FRC-monocyte axis that we have previously shown to regulate plasmablast accumulation in healthy mice (28). In this axis, stromal CCL2 in the T zone and the medulla induced local CCR2+ Ly6^hi^ monocytes to upregulate ROS expression, which then limited survival of plasmablasts that were colocalized with the CCL2-expressing FRCs and monocytes. Upon MLD of IMQ-treated CCL2-GFP reporter mice, FRCs in the draining auricular lymph nodes showed upregulated GFP, suggestive of upregulated CCL2 expression (**Figure 5A**). By anti-CCL2 staining, an increase in FRC CCL2 was also detectable in LEC^PTEN^-IMQ mice compared with control LEC^WT^-IMQ mice (**Figure 5B).** Additionally, MLD also increased FRC numbers **(Figure 5C),** which likely reflected the increased proliferation rate, as indicated by increased Ki-67 levels **(Figure 5D-E).** This expansion of FRC numbers likely added further to the level of CCL2 sensed by CCR2+ cells in the lymph node.

**Figure 5.**
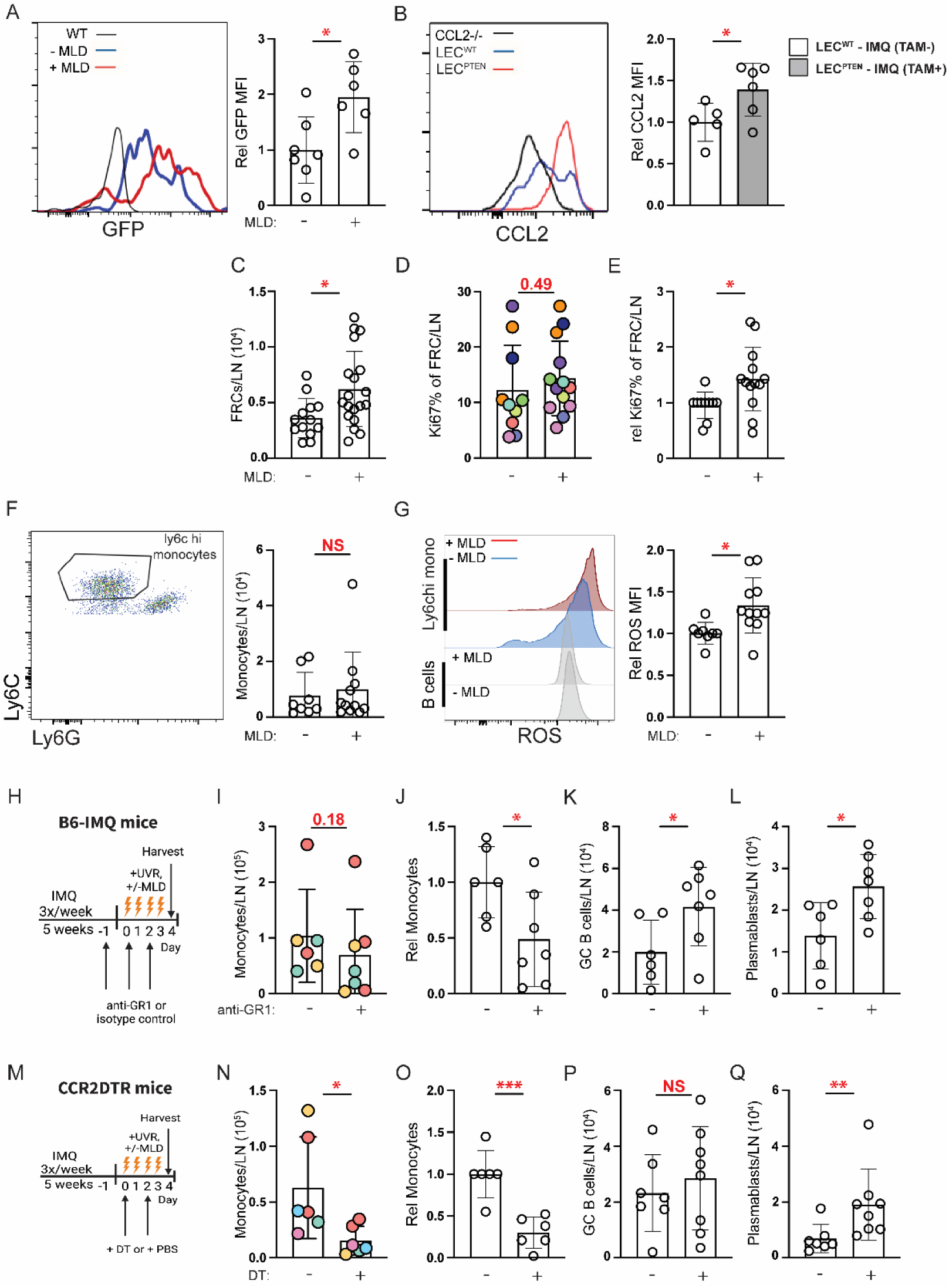
Improving lymphatic flow increases lymph node FRC proliferation, FRC CCL2, monocyte ROS generation, and limits plasmablast numbers in a monocyte-dependent manner (A-Q) Left auricular lymph nodes of indicated mice exposed to 4 days of UVR were examined. (A) FRC expression of GFP in CCL2-GFP reporter mice treated with IMQ that received MLD or control handling with UVR. (B) CCL2 expression by FRCs in LEC^PTEN^-IMQ and LEC^WT^-IMQ mice. (A, B) Representative histogram (left) and graph (right). MFI=geometric mean fluorescence intensity. (C-G) B6-IMQ mice received 4 days of +/-MLD with UVR. (C) FRC numbers. (D) Percentage of FRCs that express ki67, (E) normalized to control handled mice. (F) Ly6C^hi^ monocyte numbers, flow cytometry gating (left) and numbers (right). (G) Monocyte ROS measured using CM-H2DCFDA, representative histograms (left) and relative MFI of CM-H2DCFDA (right). (H-L) B6-IMQ mice were treated with anti-Gr-1 or isotype control at days-1, 0, and +2 of UVR and MLD treatments as shown in (H). (I) Monocyte numbers and (J) normalized to isotype control. (K) Germinal center B cell and (I) plasmablast numbers. (M-Q) CCR2-DTR mice were treated with DT at days 0 and 2 of UVR and MLD treatments as shown in (M). (N) Monocyte numbers and (O) normalized to control. (P) Germinal center B cell and (Q) plasmablast numbers. Each symbol represents one mouse; n= 6 to 19 per condition; data are from 2 (B), 3 (A, I-L), 5 (N, O), 6 (D-G, P, Q), and 7 (C) independent experiments. Normality was assessed using the Shapiro-Wilk test. If normal, unpaired t test was used. If data were not normal, Mann-Whitney U test was used. ***P<0.001, **P<0.01, *P<0.05, NS=not significant. Error bars represent SD.

Together, our results suggested that improving lymphatic flow increased the level of FRC CCL2 in draining lymph nodes.

In association with the stromal CCL2 upregulation, Ly6C^hi^ monocyte numbers were unchanged (**Figure 5F)**, but monocyte ROS expression was increased (**Figure 5G**). The monocyte ROS increase was specific to this population, as B cells did not show a similar increase (**Figure 5G**). This finding of unchanged monocyte numbers but upregulated ROS expression was consistent with the FRC-monocyte axis in healthy mice (28).

Together, these results were consistent with a model whereby improving lymphatic flow increased stromal CCL2 which then increased CCR2+ Ly6C^hi^ monocyte expression of ROS to control B cell responses in draining lymph nodes.

To test this model, we took two approaches to ask the extent to which monocytes were required for the reduced B cell responses seen with improved lymphatic flow. We used anti-Gr-1 (28) that depleted both monocytes (**Figure 5H-J**) and neutrophils (**Supplemental Figure 6A**) in B6-IMQ mice during UVR and MLD. This depletion was associated with restoration of germinal center B cell and plasmablast numbers to the higher levels seen in mice without MLD (**Figure 5K-L; compare with Figure 3L-M),** suggesting that MLD-driven reduction in B cell responses was dependent on myeloid cells. To further assess the role of monocytes without depleting neutrophils, we induced the IMQ model in CCR2-DTR mice, injected DT to deplete monocytes (**Figure 5M-O)** but not neutrophils (**Supplemental Figure 6B),** and then treated with UVR and MLD **(Figure 5M)**. Germinal center B cell numbers were not altered but plasmablast numbers were restored to the higher levels seen in mice without MLD (**Figure 5P-Q; compare with Figure 3L-M)**. The results of the Gr-1 depletion and the CCR2-DTR model together suggested that improved lymphatic flow limits plasmablast accumulation in a monocyte-dependent manner, while the reduction in germinal center B cells is mediated by other mechanisms. All together, our data suggested a model whereby restoring lymphatic flow in UVR-treated SLE mice reduces draining lymph node B cell responses at least in part by upregulating stromal CCL2 and increasing monocyte ROS to limit plasmablast accumulation.

## Discussion

Lymphatic flow is critical for clearing inflammatory mediators from peripheral tissues and communicating with draining lymph nodes, and our results suggested that the link between UVR-induced photosensitive skin responses and increased autoimmunity in SLE is at least in part due to a propensity for SLE skin to develop UVR-induced lymphatic dysfunction. By correcting this dysfunction using either MLD or a transgenic model with increased lymphatic flow, we showed that this lymphatic dysfunction contributed to both upstream cutaneous photosensitive responses and downstream draining lymph node B and T cell responses **(see Figure 6 graphical summary)**. While the contribution of lymphatic dysfunction to increasing skin inflammation is consistent with findings in non-lupus models (42, 49), we establish here that lymphatic dysfunction also contributes to B and T cell responses in lymph nodes.

Our results suggest that lymphatic dysfunction prevents optimal function of the regulatory FRC-monocyte axis in lymph nodes that normally limits plasmablast responses, and that improving lymphatic function and reducing B cell responses, over the long term, can reduce autoantibody titers. Reduced antibody titers, in turn, has the potential to lead to reduced deposition of immune complexes and limiting inflammation and damage in skin, kidneys, and other end organs. Together, our results suggested that UVR-induced lymphatic flow dysfunction is a contributing factor to lupus pathophysiology and points to lymphatic modulation of lymph node stromal function as a therapeutic target for UVR-induced disease flares.

Our finding that lymphatic flow modulates lymph node stromal phenotype in SLE models underscores the importance of the communication between tissue and lymph nodes and highlights FRC regulation as an outcome of this communication. In this setting, FRCs act as a rheostat that senses and converts peripheral tissue signals into regulators of lymph node activity. We showed that improving flow was connected functionally to the FRC CCL2-monocyte ROS axis that we had previously delineated in the setting of immune responses in non-lupus models. Consistent with the findings in healthy mice, this axis contributed to limiting plasmablast responses and not germinal center responses. Anatomically, CCL2 FRCs are positioned with plasmablasts within the T cell zone and medulla, where they are able to modulate local monocytes, and consequently the plasmasblasts (28, 50). FRCs, similar to fibroblasts in the synovium and other tissues (51, 52), are comprised of multiple subsets that have distinct functions related to their anatomic positioning within lymph nodes (25, 26); it will be interesting to further understand how different FRC subsets are modulated phenotypically and functionally by changes in lymphatic flow.

The ways by which lymphatic flow modulates FRC phenotype remains to be determined. FRCs are excellent mechanosensors and respond to environmental alterations to modulate immune function (53–55), and our results may reflect FRC sensing of changes in parameters such as shear stress as lymphatic flow changes. It is also possible that our results reflect FRC sensing of different soluble mediators originating in the skin as lymphatic flow and consequent skin inflammation is modulated (56). For example, with the MLD-induced reduction of T cell interferon gamma expression in skin, there could be reduced levels of interferon that reach the lymph node FRCs, thus altering FRC phenotype. The reduced skin inflammation could also include reduced type I IFN expression in skin, with reduced type I IFN flowing to draining lymph nodes. Here, it is interesting to note that IFNAR deletion from FRCs has been shown to upregulate FRC CCL2 expression (57), raising the possibility that reduced type I IFN coming from the skin could cause FRC CCL2 upregulation and activate the FRC-monocyte axis to limit B cell responses. Yet another potential way by which improving lymphatic flow can modulation FRC function is that while dendritic cell mobilization from skin to lymph nodes is relatively well preserved even in the face of changes in lymphatic fluid flow (58), dendritic cells can modulate FRC phenotype (59) and skin dendritic cell alterations induced by the reduced skin inflammation upon improving lymphatic flow may potentially alter FRC function upon dendritic cell migration to draining nodes. Further elucidation of how lymphatic manipulation impacts the lymph node stromal compartment will be an important future direction.

The reduced lymphatic flow and relationship to B cell responses in lupus models echoes the findings of Swartz and colleagues who showed in otherwise healthy mice that a dearth of dermal lymphatics leads over time to autoantibodies generation (17). Our data showing that lymphatic function is connected to a lymph node FRC-monocyte axis that we have previously shown in healthy mice (28) to limit B cell responses provides a potential mechanism to explain these findings. This would suggest that the lymphatic regulation of the FRC-monocyte axis is a physiologic mechanism for immune regulation, and that the effects of UVR exposure on autoimmunity in lupus is, in part, a disruption of this physiologic lymphatic-FRC-monocyte axis.

Our study leads to many more questions. It will be interesting to understand the other mechanisms by which lymphatic flow impacted lymph node immune function. For example, the lymphatic-modulated FRC-monocyte axis in lupus model mice contributed to limiting plasma cell responses and not germinal center responses. In LPR mice, reduced TFH numbers by improved lymphatic flow may have contributed to reduced germinal center responses; how improving lymphatic flow reduced TFH numbers or the T cell interferon gamma expression in both LPR and IMQ models in lymph nodes and the extent to which these changes are the consequence of FRC changes remains to be examined. It will also be interesting to understand the mechanisms by which lupus leads to lymphatic dysfunction. Type I IFN has been shown to inhibit dermal lymphatic fluid transport in the setting of a vaccinia skin infection model (60), suggesting the possibility that the high IFN-I tone in non-lesional human and murine lupus skin (39, 61–63) combined with UVR-induced IFN-I upregulation (64) could be contributing to the lymphatic dysfunction that we observe. The high interferon tone and other features of lupus skin may also contribute to lymphatic flow changes even without UVR exposure that was not captured with our Evans Blue dye assay but that may have immune consequences. Additionally, lymphatic endothelial cells play important roles in regulating the immune cells that enter and migrate within the lymphatics (65–67); the ways in which lymphatic endothelial cell phenotype is altered in lupus skin and the impact on autoimmunity will be interesting to understand going forward.

Our results have clinical implications. Potentially, MLD could be used in addition to current medical therapies to reduce local cutaneous inflammation or, over a larger area over time, to contribute to reducing systemic disease. Examination will be needed to assess the utility of MLD as an accessible and relatively inexpensive adjunct approach to ameliorate disease in SLE.

## Materials and Methods

### Study Design

The purpose of this study was to examine lymphatic function in the skin of photosensitive SLE mouse models and human SLE skin and to understand the contributions of lymphatic function on cutaneous photosensitive and lymph node responses. Laboratory mice were used as subjects for experiments. Human skin sections were also analyzed. Evans blue lymphangiography was used to assess lymphatic flow and flow cytometry was used to identify and quantify cell numbers. For experiments, sample sizes of n=3-21 animals per condition were evaluated in 1 to 11 independent experiments.

### Human Samples Staining and Imaging

FFPE sections from skin biopsies of 8 patients with SLE and/or persistently positive APL examined in (32) were used to stain for lymphatic vessels. Patients in the study had active livedo reticularis and staining for lymphatic vessels was done on biopsies taken from the purple areas of the livedo. Five patients had SLE, three of which had concomitant persistent positive APL, and three patients had persistent positive APL without SLE. Two archived samples from healthy donors were also used. Ethical approval for this study was obtained from Hospital for Special Surgery Institutional Review Board (IRB) (IRB number: 2015-256), where participants had signed written informed consents for both the initial study and for the subsequent study of skin biopsies.

Five micrometer paraffin sections were deparaffinized in xylene and rehydrated in a graded alcohol series. After a final wash with distilled water, specimen slides were placed in boiling Antigen Unmasking Solution, Tris-Based (Vector Laboratories) for 15 minutes. Slides were briefly washed in water and thereafter in phosphate buffered saline (PBS)-0.025% triton-X-100 prior to blocking non-specific binding sites with PBS-3% bovine serum albumin (BSA) for 30 minutes at room temperature. Double immunostaining was performed with 1:40 dilution in PBS/0.5% BSA of anti-PDPN (D2-40, Biolegend, San Diego, CA) and 1:100 dilution of anti-CD31/PECAM-1 (Novus Biologicals, Centennial, CO) overnight at 4°C. Slides were washed with PBS-0.025% triton-X-100 and endogenous peroxidase was inhibited by incubating slides in 3% H_2_O_2_ for 15 minutes in the dark. The sections were then washed in distilled water followed by wash buffer and incubated with alkaline phosphatase-conjugated donkey anti-mouse and horse radish peroxidase-conjugated donkey anti-rabbit secondary antibodies (both Jackson ImmunoResearch, West Grove, PA) at 1:100 dilution (prepared in tris-buffered saline (TBS) with 0.5% BSA) for 1 hour at room temperature. The reaction was revealed with a 3,3′-diaminobenzidine Substrate Kit for peroxidase and fast blue substrate (both Sigma-Aldrich, Burlington, MA) for alkaline phosphatase. At the end of incubation, slides were washed in TBS-0.025% triton-X-100, mounted with Clear-Mount (Electron Microscopy Sciences) and baked at 56°C for 20 minutes. Imaging was performed with Leica Aperio CS2 slide scanner at 40X magnification and ImageJ (NIH) analysis software used to measure the lumenal area of PDPN+CD31+ lymphatic vessels. Image analysis was performed blinded to the patient diagnosis.

### Mice

Mice between 6-15 weeks of age were used unless otherwise specified. Both male and female mice were used for experiments. All experiments were performed with age and sex matched controls. C57Bl/6, CCL2^-/-^ (68), CCL2-GFP (69), MRL^MpJ^ (MRL) and MRL-Fas^lpr^ (LPR) mice were originally from Jackson Laboratory (JAX, Bar Harbor, Maine) and bred at our facility. PTEN^fl/fl^ CreERT2 (42) were as described. CCR2-DTR (70) mice were bred at our facility. All animal procedures were performed in accordance with the regulations of the Institutional Animal Care and Use Committee at Weill Cornell Medicine (New York, NY).

### Manual lymphatic drainage

MLD was adapted to the mouse by a licensed physical therapist (CBC) with experience in MLD. MLD targeting the left ear was performed daily as specified. ProDerma smooth, powder-free latex gloves (Uniseal, Alhambra, CA) were used to minimize friction and cutaneous trauma and all movements were done at a speed of 1-2 seconds per movement. Mice were anesthetized throughout procedure. The first step was the clearing step, and this started with passive motion of the left forelimb, where the mouse was placed in a supine position, held by the paw and clockwise rotations were performed (20 repetitions) to clear the supra-clavicular and axillary area. Subsequently, stationary circular movements with light pressure of the index finger were applied on the left submandibular area followed by the auricular area (20 reps each). The mouse was then placed in the right lateral decubitus position to proceed with the reabsorption step which was performed in a specific sequence: 1. The left ear was held gently with a pincer grip (thumb and index finger) and light sweeping movements with the index finger were done on the ventral surface of the ear from distal to proximal (50 reps); 2. Placed in a prone position, the ear was gently held with a pincer grip and sweeping movements were done from distal to proximal on the dorsum of the base of the ear (where the large collecting lymphatic vessels are and can be visualized by Evan’s blue lymphangiography) (200 reps); 3. Placed back to the right lateral decubitus position, sweeping movements at the base of the ear toward the auricular lymph node were performed (20 reps); 4. Finally placed in supine position again, the same clockwise rotations of the left forelimb was performed (20 reps). Handling controls were anesthetized in the same manner as for MLD, and the mouse was placed prone and left ear was held with a pincer grip similarly to MLD for 5 minutes.

### Mouse treatments

For UVR treatments, mice were exposed to 1000 - 1500 J UVB/m^2^/day for four consecutive days using a bank of four FS40T12 sunlamps, as previously described (35). For long term treatment, mice were exposed to UVR for four consecutive days the first week and then for 3 consecutive days/week in the following weeks. To measure ear swelling after UVR exposure, a caliper (Mitutoyo, Industry, CA) was used. Each ear was measured in the anterior half of the ear three times and the average was taken.

For the IMQ-induced lupus mouse model, mice were painted on the dorsal and ventral sides of the right ear with 5% imiquimod cream (Taro Pharmaceutical) 3x/week for 4-5 weeks (∼50 mg/mouse total cumulative dose) (37).

For mice receiving tamoxifen, tamoxifen (Sigma-Aldrich) in corn oil was injected intraperitoneally (IP) at a dose of 300mg/kg/day every other day for three doses.

For monocyte depletion studies in the B6-IMQ mice, anti-Gr1 (RB6-8C5) or isotype control IgG (LTF-2) (both BioXCell, Lebanon, NH) were injected IP at a dose of 250μg in 200μl PBS on indicated days. To deplete monocytes in the CCR2-DTR mice, diphtheria toxin (Enzo Life Sciences, Farmingdale, NY) was injected IP at a dose of 250ng in 200 ul PBS.

### Evans blue lymphatic function assay

Evans blue retention assay was performed as previously described (71). Mouse ears were injected intradermally at the tip using a Hamilton syringe (#1701, 10 ul syringe) with a 30-gauge needle. 1μl of 2% Evans blue (Sigma-Aldrich) was injected and mice were euthanized 22-24 hours later. The harvested ear was placed in 300µl formamide at 58°C overnight to extract Evans blue, which was quantified by absorbance with a Multiskan Ascent plate reader (Titertek, Pforzheim, Germany) at 620nm using a titration curve.

### ROS Staining

Intracellular ROS was measured using 5-(and6)chloromethyl-2’,7’-dichlorodihydrofluorescein diacetate, acetyl ester (CM-H2DCFDA, Thermo Fisher Scientific), as in (28). The dye was reconstituted at 5 mmol/L in DMSO and stored at −20°C. Cells were prepared in RPMI and stained with 1/500 dilution of the stock in PBS for 30 minutes at 37°C prior to flow cytometric analysis.

### Flow cytometry staining and quantification

For flow cytometric staining of skin, single cell suspensions were generated as previously described (72). In brief, ears were finely minced, digested in type II collagenase (616 U/mL; Worthington Biochemical Corporation, Lakewood, NY), dispase (2.42 U/mL; Life Technologies, Carlsbad, CA), and DNAse1 (80 μg/mL; Sigma-Aldrich), incubated at 37°C while shaking at 100 rpm, triturated with glass pipettes, and filtered. For staining of lymph node cells, hematopoietic cells from lymph nodes were obtained by mashing the lymph nodes and extruding the cells through a 70μm strainer. Stromal cells were obtained as previously described (73); lymph nodes were minced, digested with type II collagenase (616 U/mL) and DNAase1 (40μg/mL) at 37°C while shaking at 50 rpm, triturated with glass pipettes, and filtered.

To count cells, the single cell suspension from the whole lymph node or ear was washed and resuspended in 300 ul of buffer. Ten ul was taken to be counted on the Multisizer 4e Coulter Counter (Beckman Coulter) and this count was used to calculate total number of cells per lymph node. One to two million cells per sample were stained, and most of the sample was run on the flow cytometer. To obtain absolute number of a particular population of cells per ear or lymph node, the frequency of these cells (of the total) in the facs analysis was multiplied by the total number of cells per tissue (as calculated by the Counter Counter data).

For flow cytometry analysis, gating of specific populations was performed after excluding debris, doublets, and dead cells using 4’,6-diamidino-2-phenylindole dihydrochloride (DAPI, Invitrogen) for non-fixed cells. Antibodies are from Biolegend unless otherwise specified. Samples were treated with anti-mouse CD16/32 (Fc block, clone 93) prior to staining with additional antibodies. Gating strategies and antibodies used--B cells: CD45+(30-F11) B220+(RA3-6132); monocytes: CD45+, B220-, CD3-(145-2C11),

CD11b+(M1/70), Ly6C^hi^(HK1.4), Ly6G-(1A8); neutrophils: CD45+, CD11b+, Ly6C^med^, Ly6G+; germinal center B cells: CD3-, B220+, GL7+(GL7), PNA+(Vector Laboratories); plasmablasts: CD3-, B220med-lo, CD138+(281-2). In digested tissues, plasmablasts were identified by either intracellular IgG (IgG1-A85-1, IgG2a/b-R2-40, IgG3-R40-82, all BD Biosciences) or intracellular Ig kappa+(187.1) (Southern Biotech) using BD Cytofix/Cytoperm kit (BD Biosciences) in lieu of CD138. Plasmablasts were confirmed by Ki67(16A8) staining. Lymphatic endothelial cells: CD45-, CD31+ (390), PDPN+ (8.1.1); blood endothelial cells: CD45-CD31-, PDPN-; fibroblastic reticular cells: CD45-, CD31-, PDPN+. CD4 T cells: CD45+CD4+TCRab+ (H57-597); regulatory T cells: TCRab+, CD4+, CD25+ (PC61), intracellular Foxp3+ (FJK-16s); follicular helper T cells: TCRab+, CD4+, CXCR5+ (L138D7), PD1+ (29F.1A12); Th1 cells: TCRab+, CD4+, Foxp3-, CXCR5-, intracellular IFNg+ (XMG1.2, eBioscience); Th17 cells: TCRab+, CD4+, Foxp3-, CXCR5-, intracellular IL17+ (TC11-18H10, BD Biosciences). The intracellular stains for CD4+ T cell subset was done after fixing cells with the eBioscience™ Intracellular Fixation & Permeabilization Buffer Set (Cat: 88-8824-00). For Th1 and Th17 cells, staining was done after cells were stimulated with Cell Activation Cocktail with Brefeldin A ( Biolegend, Cat no. 423304) in 37C incubator for 4 hours. CCL2 was identified by either using CCL2-GFP reporter mice or by staining. Staining for CCL2 was performed using CCL2-FITC (245, Invitrogen) and signal was amplified by staining with anti-FITC-biotin (1F8-1E4, Jackson ImmunoResearch) and subsequently streptavidin-FITC (Invitrogen). A negative control for GFP signal was done by using non-transgenic mice in CCL2-GFP mice experiments. CCL2^-/-^ mice were used as negative control for staining experiments.

For flow cytometry analysis, cells were analyzed using a FACSCanto or FACSSymphony (BD Biosciences) and FlowJo Software (Tree Star).

### Cell Sorting

For qPCR of skin cell populations, cells from ear skin were pooled and then sorted using a BD Influx (BD Biosciences). LECs were selected as DAPI-CD45-PDPN+CD31+ cells. BECs were selected as DAPI-CD45-PDPN-CD31+ cells. Fibroblasts were selected as DAPI-CD45-PDPN+CD31-cells. Macrophages were selected as DAPI-CD45+CD11b+F4/80+ cells.

### RNA Extraction

RNA was extracted from sorted cells using an RNAEasy Plus Kit (Qiagen, Germantown, MD) and quality confirmed on a BioAnalyzer 2100 (Agilent Technologies, Santa Clara, CA).

### Real time PCR

cDNA was synthesized (iScript kit, Bio-Rad, Hercules, CA) from extracted RNA and real-time PCR (iQ SYBR-Green Supermix kit, Bio-Rad)) was performed using primers for PTEN (Mm_Pten_1_SG QuantiTect Primer Assay, Qiagen) and GAPDH (R: TTGAAGTCGCAGGAGACAACCT, F: ATGTGTCCGTCGTGGATCTGA).

### Autoantigen Microarray

Blood was collected from mice, left at room temperature for 1 hour, and then centrifuged at 3000rpm for 3 minutes. The supernatant containing the serum was then collected and frozen in-80C until ready to be shipped for autoantigen microarray profiling at the Genomics and Microarray Core Facitlity, UT Southwestern Medical Center. At the core, the samples are treated with DNAse I, diluted 1:50, and incubated with autoantigen array. The autoantibodies binding to the antigens on the array are detected with Cy3 labeled anti-IgG and cy5 labeled anti-IgM and the arrays are scanned with GenePix® 4400A Microarray Scanner. The images are analyzed using GenePix 7.0 software to generate GPR files. The averaged net fluorescent intensity (NFI) of each autoantigen is normalized to internal controls (IgG or IgM).

### Statistics

For figures showing normalized values, each individual replicate experiment was normalized for that experiment. For experiments that contained more than one control sample, the mean was obtained for the control samples, and the individual control and experimental samples were divided by this mean value to normalize to the control mean. The Shapiro-Wilk test was used to test for normality. Unpaired t test was used for normal data and Mann-Whitney U test was used otherwise.

## Author Contributions

WGA, NS, and TTL conceived the study; MJH, WGA, MLC, AR, ESS, JHS, JS, NS, WDS, DCD, CBC, RPK, YC designed and performed the experiments and analyzed the data; ES, DE, SS, RPK, and BM provided key reagents and intellectual input. CBC and SAR designed experiments and provided intellectual input. TTL designed, supervised, and interpreted experiments. MJH, WGA, and TTL wrote the manuscript; all authors contributed to critical reading and editing.

## Acknowledgments

The authors thank the Lu Lab for critical reading of the manuscript. This work was supported by R01AI079178-121S1 (MJH), NIH T32AR071302-01 to the Hospital for Special Surgery Research Institute Rheumatology Training Program (NS and WDS), NIH MSTP T32GM007739 to the Weill Cornell/Rockefeller/Sloan-Kettering Tri-Institutional MD-PhD Program (WDS), Emerson Collective Cancer Research Fund 691032 (RPK), NIH R01AI079178 (TTL), R01AR084694 (TTL), R21 AR081493 (TTL), DOD W81XWH-22-1-0627(TTL), Lupus Research Alliance (TTL), the St. Giles Foundation (TTL); and NIH Office of the Director grant S10OD019986 to Hospital for Special Surgery.

