## Supplemental Data for "Lymphatic dysfunction in lupus contributes to cutaneous photosensitivity and lymph node B cell responses"

Supplemental Figures 1-6 for Howlader, Ambler et al.

### Lymphatic dysfunction in lupus contributes to cutaneous photosensitivity and lymph node B cell responses

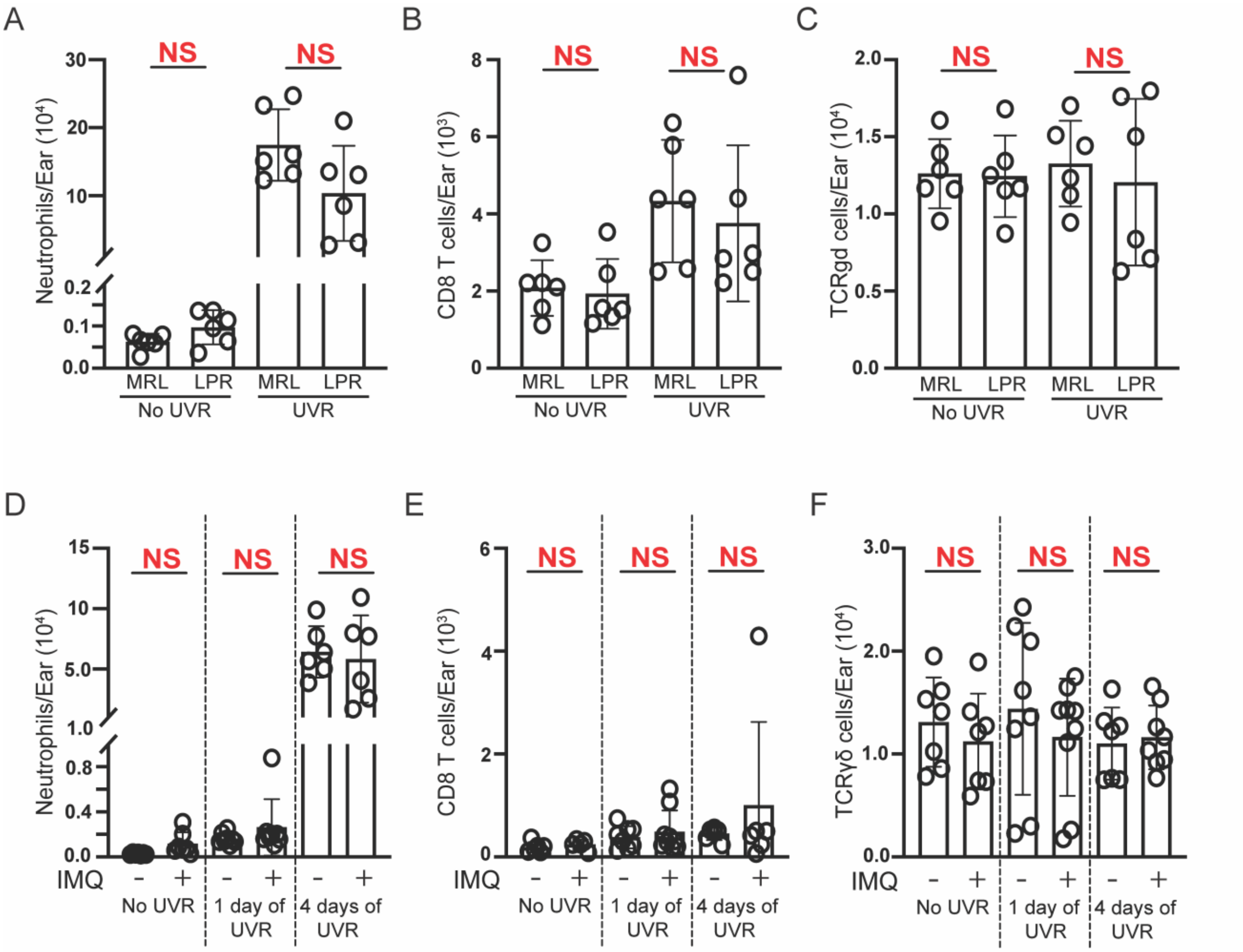


**Supplemental Figure 1. LPR and B6-IMQ mice show no changes in neutrophil, CD8 T cell, and TCRγδ T cell numbers in left ear after UVR exposure**

(A-C) LPR mice with no UVR or treated with 4 days UVR had left ears collected 24 hours after final UVR dose. (A) Neutrophil, (B) TCRαβ CD8 T cell, (C) TCRγδ T cell numbers. (D-F) B6-IMQ mice were treated with zero, 1 day, or 4 days of UVR prior to left ear harvest. (D) Neutrophil, (E) TCRαβ CD8 T cell, and (F) TCRγδ cell numbers. Each symbol represents one mouse; n= 5 to 9 per condition; data are from 2 (A-C), 8 (E, F), and 11 (D) independent experiments. The Shapiro-Wilk test was used to test for normality. Unpaired t test was used for normal data and Mann-Whitney U test was used otherwise. ***P<0.001, **P<0.01, *P<0.05, NS=not significant. Error bars represent SD.

A

| Apoptosis pathways that are different between IMQ painted and unpainted ears | FDR | log2 FC |
|---|---|---|
| REACTOME_APOPTOTIC_CLEAVAGE_OF_CELL_ADHESION_PROTEINS | 4.16E-05 | 0.44 |
| REACTOME_DEFECTIVE_INTRINSIC_PATHWAY_FOR_APOPTOSIS | 4.99E-05 | 0.30 |
| REACTOME_APOPTOTIC_EXECUTION_PHASE | 1.92E-04 | 0.15 |
| REACTOME_APOPTOTIC_CLEAVAGE_OF_CELLULAR_PROTEINS | 2.63E-03 | 0.12 |
| REACTOME_APOPTOSIS | 2.64E-03 | 0.10 |
| REACTOME_CYTOCHROME_C_MEDIATED_APOPTOTIC_RESPONSE | 2.16E-02 | 0.08 |
| REACTOME_INTRINSIC_PATHWAY_FOR_APOPTOSIS | 2.87E-02 | 0.07 |

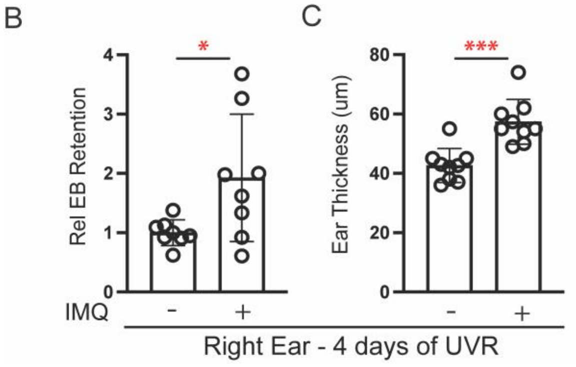


**Supplemental Figure 2. Direct IMQ exposure of the painted right ear in B6-IMQ mice upregulates damage pathways, and UVR exposure results in impaired lymphatic flow and increased swelling of right ear**

Tissue apoptotic pathways are upregulated in IMQ-painted (right) ear compared to unpainted (left) ear in the B6-IMQ model. Bulk RNA seq data of IMQ painted and unpainted ear skin were previously published (1). This dataset was queried for significantly different expression levels of apoptotic pathways in the Reactome Pathway Database, and the results are shown. (B-C) B6-IMQ mice were treated with 4 days of UVR and painted right ear was assessed for (B) Evans blue dye retention and (C) ear thickness. Each symbol represents one mouse; n= 8 to 9 per condition; data are from 4 (B) and 6 (C) independent experiments. The Shapiro-Wilk test was used to test for normality. Unpaired t test was used for normal data and Mann-Whitney U test was used otherwise. ***P<0.001, **P<0.01, *P<0.05, NS=not significant. Error bars represent SD.

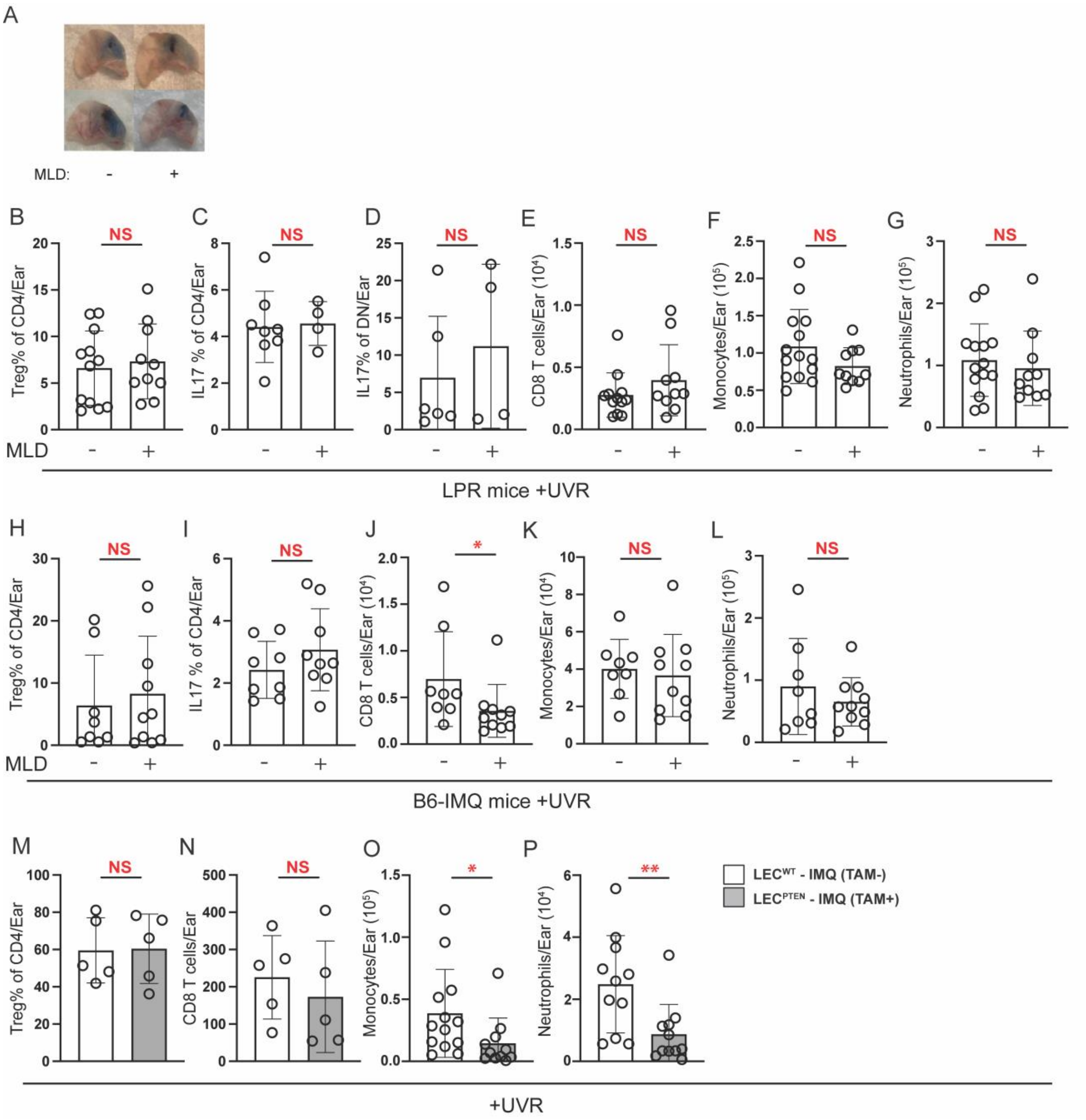


**Supplemental Figure 3. Additional assessment of immune cell changes in skin with improved lymphatic flow**

(A) Ears illustrating visible changes in Evans blue dye retention with MLD. Left ears of LPR mice that received 4 days of UVR +/- MLD were injected with Evans blue dye at 1 day after final UVR dose and ear harvested 1 day later. Left ears of (B-G) LPR and (H-

L) B6-IMQ mice that received UVR exposure with or without MLD for 4 days were examined by flow cytometry. (B, H) Percentage of TCR$\alpha\beta$ CD4 T cells that were Foxp3+CD25+ Tregs and (C, I) IL-17+ Th17 cells. (D) Percentage of DN cells expressing IL17. (E, J) CD8 T cell, (F, K) monocyte, and (G, L) neutrophil numbers. (M-P) Left ears of $LEC^{PTEN}$-IMQ and $LEC^{WT}$-IMQ mice that received 4 days of UVR exposure were examined similarly to above for (M) Treg percentage of CD4 T cells and (N) CD8 T cell, (O) monocyte, and (P) neutrophil numbers. Each symbol represents one mouse; n= 4 to 14 per condition; data are from 2 (C, D, M, N), 3 (H-L), 4 (B, E-G), and 5 (O, P) independent experiments. The Shapiro-Wilk test was used to test for normality. Unpaired t test was used for normal data and Mann-Whitney U test was used otherwise. ***P<0.001, **P<0.01, *P<0.05, NS=not significant. Error bars represent SD.

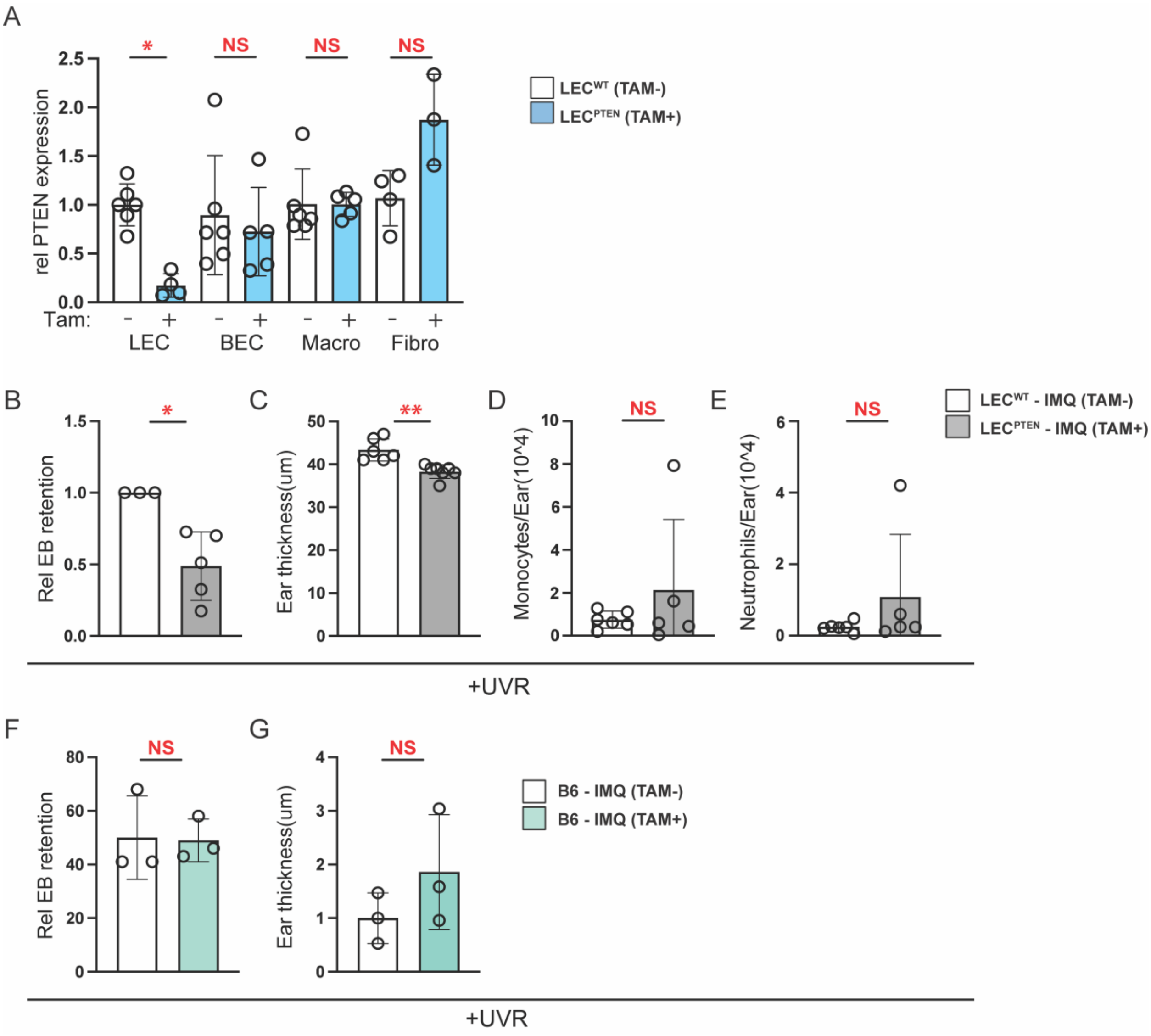


**Supplemental Figure 4. Tamoxifen-treated Flt4Cre$^{ERT2}$PTEN$^{fl/fl}$ mice had selective knockout of PTEN in LECs and right ear assessment in LEC$^{PTEN}$-IMQ mice and associated controls**

(A) PTEN expression as determined by qPCR of lymphatic endothelial cells (LECs), blood endothelial cells (BECs), macrophages, and fibroblasts sorted from ear skin of Flt4Cre$^{ERT2}$PTEN$^{fl/fl}$ mice +/- tamoxifen. (B-G) LEC$^{PTEN}$-IMQ and LEC$^{WT}$-IMQ mice (B-E) and B6-IMQ mice +/- tamoxifen (F, G) were treated with 4 days of UVR and IMQ-painted right ears were assessed for Evans Blue retention (B, F), ear thickness (C, G), monocytes (D) and neutrophils (E). Each symbol represents one mouse; n= 3 to 7 per condition; data are from 2 (D-G), 3 (A, B), and 4 (C) independent experiments. The Shapiro-Wilk test was used to test for normality. Unpaired t test was used for normal data and Mann-Whitney U test was used otherwise. ***P<0.001, **P<0.01, *P<0.05, NS=not significant. Error bars represent SD.

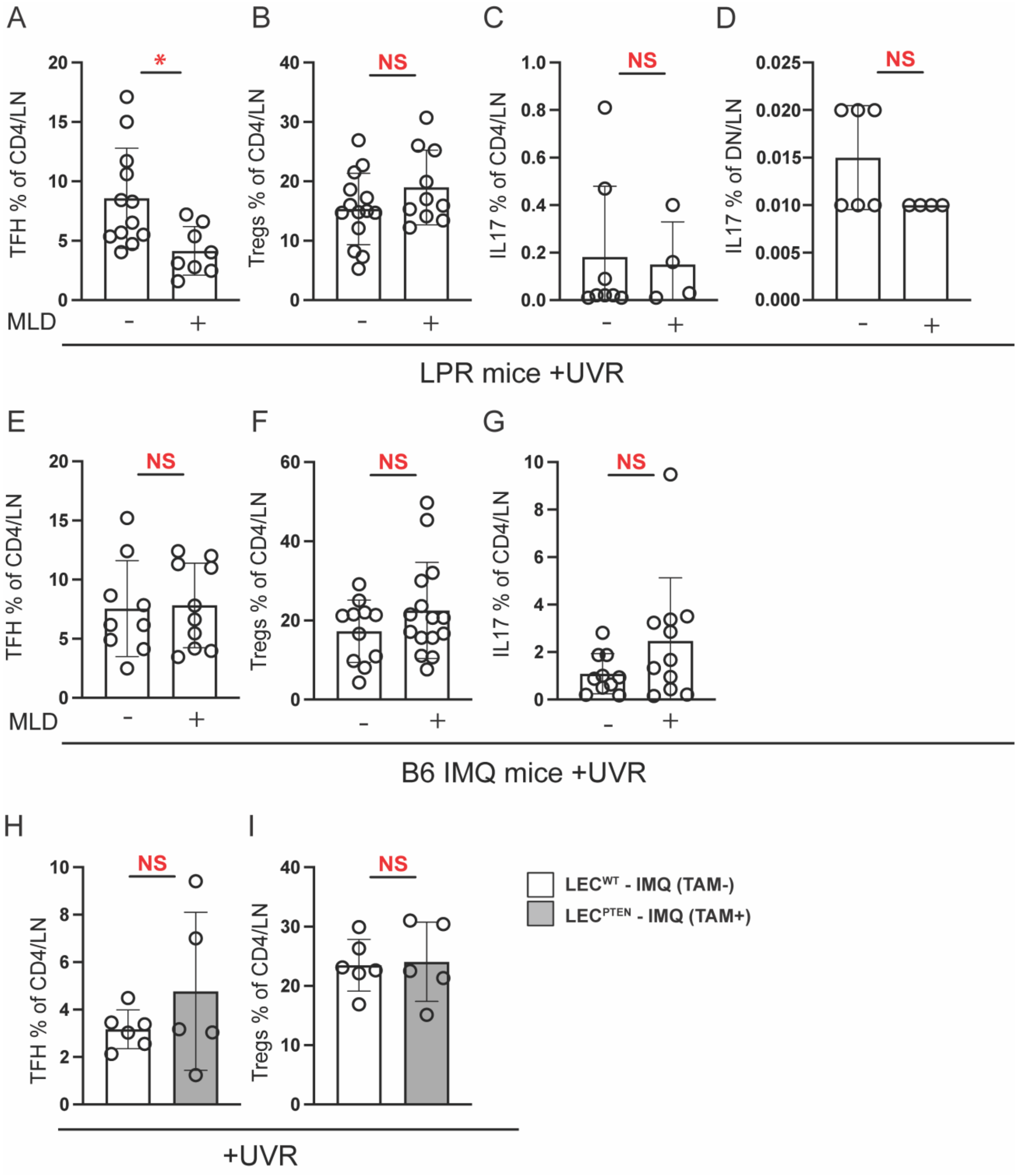


**Supplemental Figure 5. Additional assessment of immune cell changes in draining lymph nodes with improved lymphatic flow**

(A-G) Left auricular nodes of LPR (A-D) and B6-IMQ (E-G) mice that received UVR +/- MLD for 4 days were examined by flow cytometry. (A,E) T follicular helper, (B,F) Treg, and (C, G) Th17 cells, as a percentage of TCRαβ+CD4+ T cells. (D) Percentage of TCRαβ+CD3+CD4-CD8- DN T cells that express IL-17. (H-I) Left auricular lymph nodes of $LEC^{PTEN}$-IMQ and $LEC^{WT}$-IMQ mice were examined for (H) TFH and (I) Treg cells as a percentage of CD4 T cells. Each symbol represents one mouse; n= 4 to 15 per condition; data are from 2 (C, D, H, I), 3 (A, E, G), and 4 (B, F) independent experiments. The Shapiro-Wilk test was used to test for normality. Unpaired t test was used for normal data and Mann-Whitney U test was used otherwise. ***P<0.001, **P<0.01, *P<0.05, NS=not significant. Error bars represent SD.

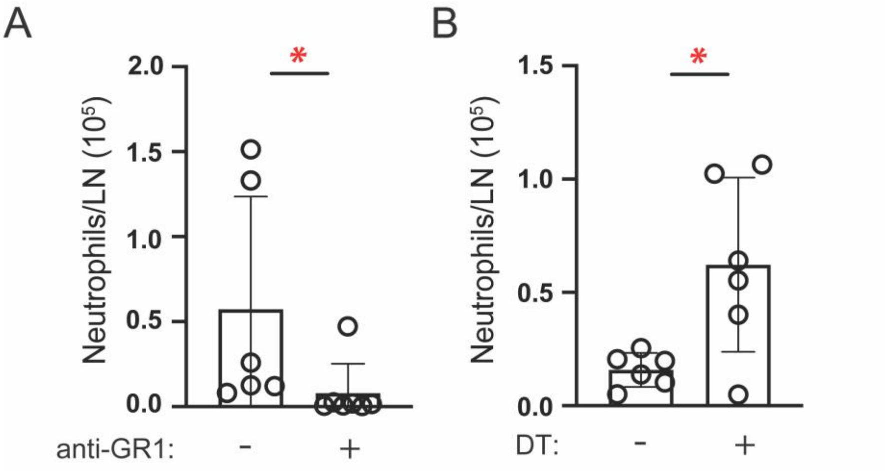


**Supplemental Figure 6. Neutrophils are depleted with anti-Gr-1 and not depleted with DT treatment of CCR2-DTR mice.**

(A) B6-IMQ mice were treated with anti-Gr-1 or isotype control at days -1, 0, and +2 of UVR and MLD treatments and left auricular lymph node was examined for CD11b+Ly6C+Ly6G+ neutrophil numbers. (B) CCR2-DTR mice painted with IMQ to induce the IMQ model were treated with DT at days 0, and +2 of UVR and MLD treatments, and left auricular lymph node was examined for neutrophil numbers. Each symbol represents one mouse; n= 6-7 per condition; data are from 3 (A) and 5 (B) independent experiments. The Shapiro-Wilk test was used to test for normality. Unpaired t test was used for normal data and Mann-Whitney U test was used otherwise. ***P<0.001, **P<0.01, *P<0.05, NS=not significant. Error bars represent SD.
